# Structural Basis of SARS-CoV-2 Spike Protein Priming by TMPRSS2

**DOI:** 10.1101/2020.04.21.052639

**Authors:** Mushtaq Hussain, Nusrat Jabeen, Anusha Amanullah, Ayesha Ashraf Baig, Basma Aziz, Sanya Shabbir, Fozia Raza

## Abstract

Entry of SARS-CoV-2, etiological agent of COVID-19, in the host cell is driven by the interaction of its spike protein with human ACE2 receptor and a serine protease, TMPRSS2. Although complex between SARS-CoV-2 spike protein and ACE2 has been structurally resolved, the molecular details of the SARS-CoV-2 and TMPRSS2 complex are still elusive. TMPRSS2 is responsible for priming of the viral spike protein that entails cleavage of the spike protein at two potential sites, Arg685/Ser686 and Arg815/Ser816. The present study aims to investigate the conformational details of complex between TMPRSS2 and SARS-CoV-2 spike protein, in order to discern the finer details of the priming of viral spike and to point candidate drug targets. Briefly, full length structural model of TMPRSS2 was developed and docked against the resolved structure of SARS-CoV-2 spike protein with directional restraints of both cleavage sites. The docking simulations showed that TMPRSS2 interacts with the two different loops of SARS-CoV-2 spike protein, each containing different cleavage sites. Key functional residues of TMPRSS2 (His296, Ser441 and Ser460) were found to interact with immediate flanking residues of cleavage sites of SARS-CoV-2 spike protein. Compared to the N-terminal cleavage site (Arg685/Ser686), TMPRSS2 region that interact with C-terminal cleavage site (Arg815/Ser816) of the SARS-CoV-2 spike protein was predicted as relatively more druggable. In summary, the present study provide structural characteristics of molecular complex between human TMPRSS2 and SARS-CoV-2 spike protein and points to the candidate drug targets that could further be exploited to direct structure base drug designing.

## Introduction

The recent pandemic of COVID-19 is the third outbreak of the diseases caused by beta coronavirus in humans, following Severe Acute Respiratory Syndrome (SARS) and Middle Eastern Respiratory Syndrome (MERS) (Cui et al., 2019). By 13th March 2020, over 1.7 million of global population has been infected with mortality rate of 21% in closed cases (WHO COVID-19 situation report-84). Genetically, etiological agent of COVID-19, SARS-CoV-2, is closely related to SARS-CoV compared to MERS-CoV (Wu et al., 2020). Similarly, like for SARS-CoV, Angiotensin Converting Enzyme-2 (ACE2) has been identified as the primary receptor for SARS-CoV-2 spike protein (Li, 2015; Lan et al., 2020). Whereas MERS-CoV spike protein interacts with the DiPeptidyl Peptidase 4 (DPP4) as the first site of attachment to the host cell (Li, 2015). Spike protein of SARS-CoV-2 is 1273 amino acid long protein with two functionally distinct regions, S1 and S2, involved in the attachment and entry of the virus, respectively. SARS-CoV-2 entry in the host cell is mediated by proteolytic cleavage of its spike protein, a process dubbed as priming. Recently, human Transmembrane Protease Serine 2 (TMPRSS2) has been shown to carry out the priming of the SARS-CoV-2 spike protein by generating two distinct fragments of the viral spike protein, S1/S2 and S2’ (Hoffman et al., 2020).

Recently, co-crystal structure of SARS-CoV-2 spike protein complexed with ACE2 receptor has been resolved unraveling the finer details of intermolecular interactions (Lan et al., 2020). The ACE2-SARS-CoV-2 complexes not only indicate the potential of cross species transmission but also open a window for designing and/or screening of disruptor molecules that could potentially inhibit the attachment of the virus with the host cells (Lan et al., 2020; Ortega et al., 2020). However, no complex structure of SARS-CoV-2 spike protein with TMPRSS2 has been resolved to date. Moreover, the molecular structure of human TMPRSS2 protein is also not known. Resultantly, structural details of intermolecular interactions between SARS-CoV-2 and TMPRSS2 are largely unknown. Although, like many other protease inhibitors (Zhang et al., 2020), TMPRSS2 inhibitor has been suggested and/or shown to antagonize the entry of the virus into the host cells (Hoffman et al., 2020). This study aims to investigate the interaction points between TMPRSS2 and SARS-CoV-2 spike protein using an array of bioinformatic tool. The findings not only provide structure-function relationship of TMPRSS2 of humans but also predict the sites of interactions between TMPRSS2 and SARS-CoV-2 spike protein. This could lead to the development and/or directed screening of disruptor and/or inhibitor molecules.

## Methodology

### Data mining for structures

Protein sequence of human TMPRSS2 (O15393) was retrieved from UniProt and subjected to PDB Blast to identify the homologous structure on the basis of query coverage and sequence identity. Atomic coordinates of SARS-CoV-2 spike protein (PDBid: 6VSB) and Hepsin (PDBid: 1Z8G) were retrieved from RCSB protein data bank (Wrapp et al., 2020; Herter et al., 2005; GoodSell et al., 2020). Probability of the protein crystallization for TMPRSS2 was predicted using XtalPred server (Slabinski et al., 2007).

### Molecular modelling

N-terminal region (1-148) of TMPRSS2 including LDL-receptor class A domain was modeled using I-TASSER (Roy et al., 2010) due to the unavailability of template with sufficient homology. Since, hepsin molecule was found to share noticeable homology with the SRCR and peptidase S1 domains of TMPRSS2, a full length model of TMPRSS2 were later developed using Modeller 9.16 (Webb and Sali, 2016) taking model developed by I-TASSER and hepsin (PDBid: 1Z8G) as templates. Full length model of TMPRSS2 was further refined for Gibb’s free energy and conformation of the loops. The structural quality of the model was assessed by MolProbity for Ramachandran Plot (Chen et al., 2010) and ProSA (Wiederstein and Sippl, 2007). Finally, the full length model was superimposed over template (PDBid: 1Z8G) and root mean square deviation in carbon alpha back bone was measured in Å using Swiss PDB Viewer v4.1.0 (Johansson et al., 2012).

### Molecular docking

HADDOCK 2.2 webserver (Van Zundert et al., 2016) was used to conduct molecular docking between SARS-CoV-2 spike protein and TMPRSS2. The input include atomic coordinates of SARS-CoV-2 spike protein (PDBid: 6VSB) and constructed full length molecular model of TMPRSS2. Two separate docking simulations were run for each cleavage site of the viral spike protein. Reported cleavage sites (Hoffman et al. 2020) on spike protein were defined as active residues for SARS-CoV-2, whereas substrate binding sites and catalytically active sites were recognized as active residues of TMPRSS2. HADDOCK congregated all docking simulations into clusters and ranked them according to the HADDOCK score which is the function of linear combination of Van der Waals energy, electrostatic energy, desolvation energy, restraint violation energy and buried surface area. The cluster with least HADDOCK score was selected for further assessment. Binding affinity and different types of interactions like charged-charged, charged-polar, charged-apolar, polar-polar, polar-apolar and apolar-apolar were identified using PRODIGY webserver (Xue et al., 2016). Interacting residues of TMPRSS2 for SARS-CoV-2 spike protein were separated and dimensions and druggability were assessed using DoGSiteScorer (Volkamer et al., 2012). All structures were visualized using DS visualizer 2016.

## Results

### Molecular model of TMPRSS2

Human TMPRSS2 is 492 amino acid long protein with three functional domains: an N-terminal LDL-receptor class A domain (113-148) followed by SRCR (153-246) and finally at C-terminal peptidase S1 domain spanning from 256 to 487 amino acid (Figure 1A). Till now molecular structure of the protein has not been resolved and our XtalPred analysis showed the least possibility for this molecule to be crystalized, potentially due to the high percentage of coiled structure, isoelectric point and surface hydrophobicity (Figure 1B). This may be the reason that since the first report of TMPRSS2 in year 1997 (Paoloni-Giacobino et al., 1997), the structure has not been resolved by X-ray crystallography. Nevertheless, computational based molecular modelling approaches have evolved since then and come of age in terms of accuracy and reliability with new tools and server being available (Roy et al., 2010; Webb and Sali, 2016). Therefore, we used multiple approaches to develop the full length molecular model of TMPRSS2. The finally selected refined model of TMPRSS2 has 96.32% residues within the allowed regions of Ramachandran plot, which is acceptable considering the N-terminal portion of the protein was predicted to be intrinsically disordered. Secondly, it has been demonstrated rather frequently that many of the resolved structures of the proteins such as USP7 (PDB id: 2F1Z) have more than 20% of the residues outside the allowed region in Ramachandran plot. Moreover, Gibbs Free energy values (−14212.818 KK/mol) and ProSA Z score (−6.89) both suggest structural reliability of the model (Figure 1C;D).

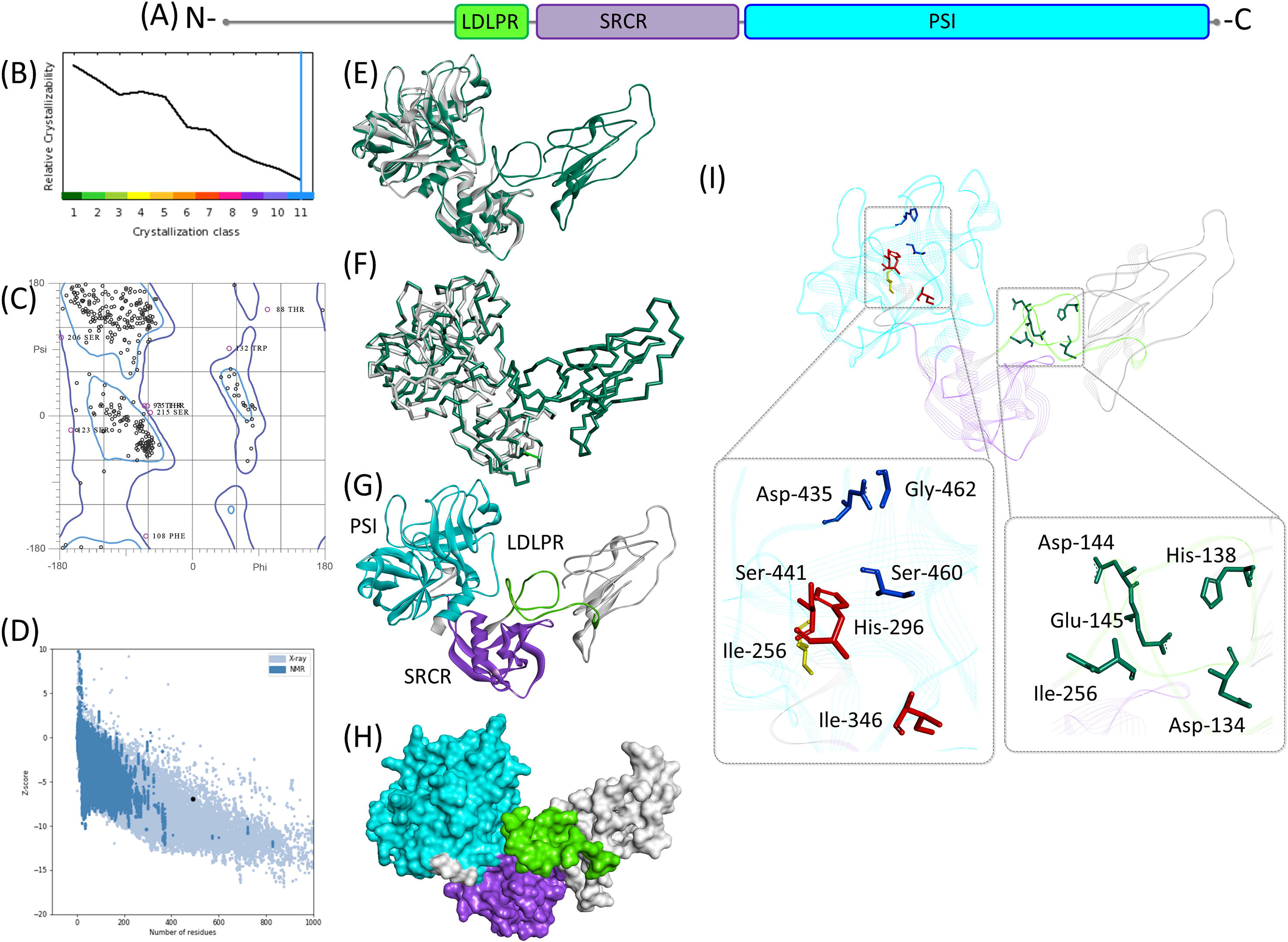

### Molecular structure of TMPRSS2

Full length molecular models of TMRPSS2 has considerable structural homology with the template molecule (PDB: 1Z8G), where the deviation between the Cα back bone of model and template was found as 0.33Å (Figure 1E;F). All three domains, LDL-receptor class A, SRCR and peptidase S1, formed distinct structural units in the molecular model. N-terminal region and LDL-receptor class A of TMPRSS2 were found more or less unstructured (Figure 1G;H). Putative Ca^2+^ biding residues (Asp134, His138, Asp144, Glu145 and Ile256) were found on a loop linking N-terminal of the protein with SRCR domain. Structurally, SRCR domain comprises an α helix and multiple anti parallel β sheets, potentially stabilized by two disulfide bonds between Cys172-Cys231 and Cys185-Cys241 (Figure 1I). Overall structural conformation of the domain showed uncanny resemblance with the SRCR domain found in MARCO receptor and hepsin (Herter et al., 2005; Ojala et al., 2007). The C-terminal of TMPRSS2 has a large catalytic domain with typical structural features of chymotrypsin family serine proteases (Mönttinen et al., 2019). The triad of catalytically active site residues (His296, Ile346 and Ser441) and substrate binding sites (Asp435, Ser460 and Gly462) were found sandwiched between two six stranded β barrels of nearly equal size (Figure 1I). The inter-residual distance between the catalytically active residues ranges from 7.965Å to 10.263Å (Supplementary figure). In comparison, the inter-residual distance between substrate binding sites ranges from 7.409Å to 11.765Å (Supplementary figure). The globular conformation of the domain is likely be stabilized by four disulfide bonds between Cys244-Cys365, Cys281-Cys297, Cys410-Cys426 and Cys437-Cys465.

### Interaction of TMPRSS2 with SARS-CoV-2 Spike protein

The entry of the SARS-CoV-2 in the host cell is driven by the proteolytic cleavage of its spike protein resulting in the formation of two fragments, S1/S2 and S2’ (Hoffman et al., 2020). The precise positioning of the proteolytic cleavage sites have been mapped by sequence comparison and found to be at the junction of Arg685/Ser686 and Arg815/Ser816. The cleavage at the later site results in the production of S1/S2 and S2’ fragments, which is necessary for the viral entry into the cells. This provides an excellent basis on which docking simulations could be directed. Thereby, in this study we run an independent docking simulation for each site. The conformation of the complex between TMRPSS2 and SARS-CoV-2 in selected docking pose revealed that both cleavage sites of SARS-CoV-2 spike protein are present at the flexible loops and interacts with one of the β barrel of the catalytic domain of TMPRSS2 (Figure 2A-C; 3A-C). At the first cleavage site (Arg685/Ser686) of the spike protein, His296 of TMPRSS2 formed a hydrogen bond and electrostatic interaction with Arg682 of the spike protein (Table 1; Figure 2D). Whereas, at the second cleavage site (Arg815/Ser816), out of the three residues of catalytic triad, His296 and Ser441 established hydrogen bond interactions with Pro809, Lys814 and Ser810 of the SARS-CoV-2 spike protein (Table 1; Figure 3D). Ser810 also formed a hydrogen bond and hydrophobic interaction with Ser460, substrate binding site, and His296, catalytic site of TMPRSS2, respectively (Table 1; Figure 3D). Since the functionally important residues of TMPRSS2 interact with the amino acids that immediately flank the cleavage site, this raises a possibility that upon interaction with the viral spike protein, the later may undergo conformational changes that may bring Arg685/Ser686 and Arg815/Ser816 of the SARS-CoV-2 spike protein in line with the active site cleft of TMPRSS2. Nevertheless, Ser441 of TMPRSS2, that has been demonstrated as the most critical residue for the proteolytic cleavage of viral spike protein (Böttcher et al., 2006; Shirogane et al., 2008), were found interacting with several flanking residue of cleavage site found in SARS-CoV-2 spike protein (Table 1; Figure 3D). This represent the importance of neigbouring residues in the establishment of molecular complex between TMPRSS2 and SARS-CoV-2 spike protein. An unpublished study (DOI: 10.1101/2020.02.08.926006) with partial modelled structure of TMPRSS2 also corroborate our findings. Furthermore, involvement of charged residues in most of the intermolecular interactions and binding affinity values (−13.8 Kcal/mol) represent the reliability of the complex in terms of structural conformation (Table 1). In order to explore the druggability, interacting residues of TMPRSS2 for each cleavage site of SARS-CoV-2 spike protein were assessed for the volume, area and drug score. Consistent to the molecular docking simulations, intermolecular interactions between TMPRSS2 and Arg815/Ser816 of SARS-CoV-2 spike protein appeared as a suitable target for drug designing and development (Figure 2E-F; 3E-F).

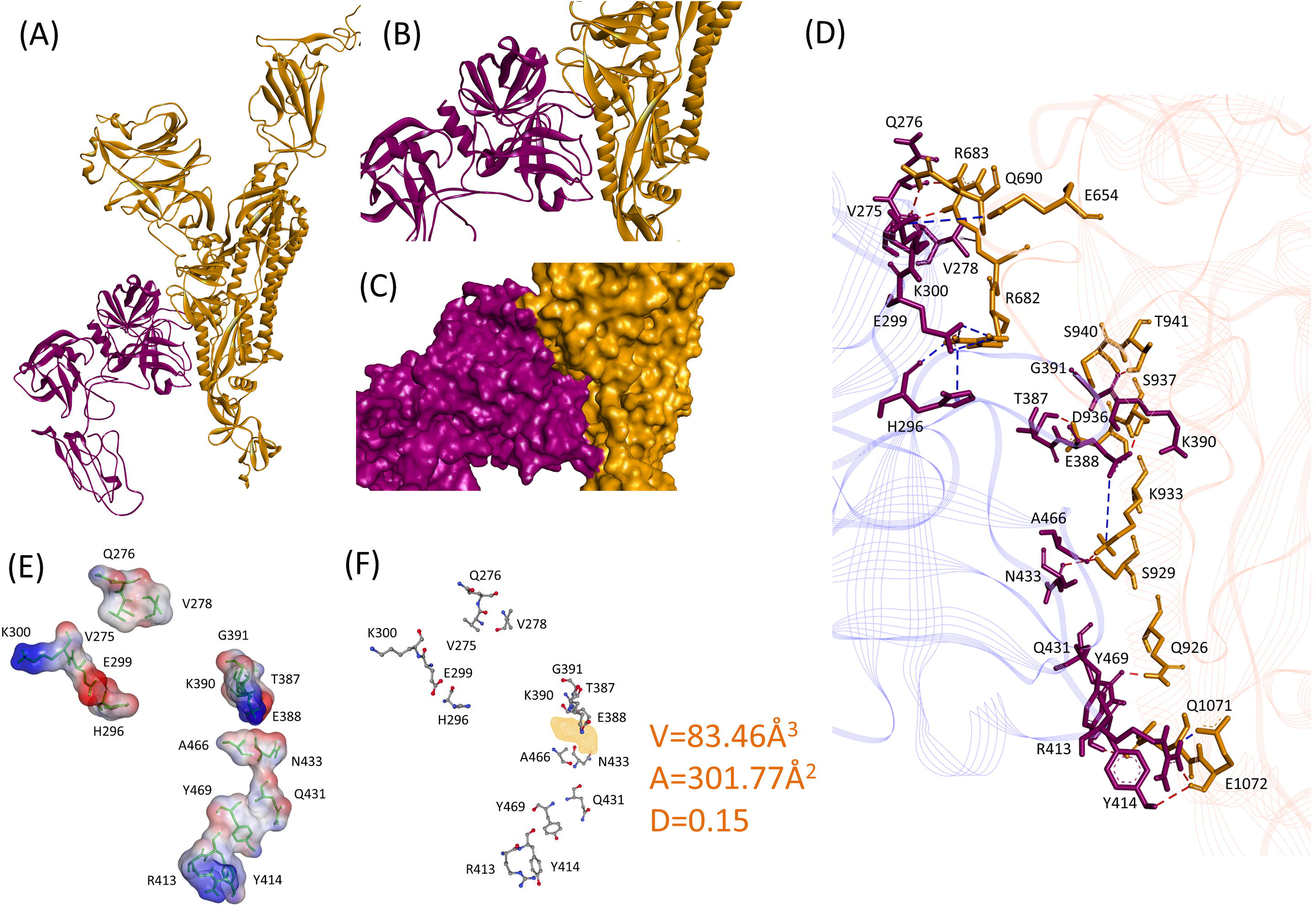

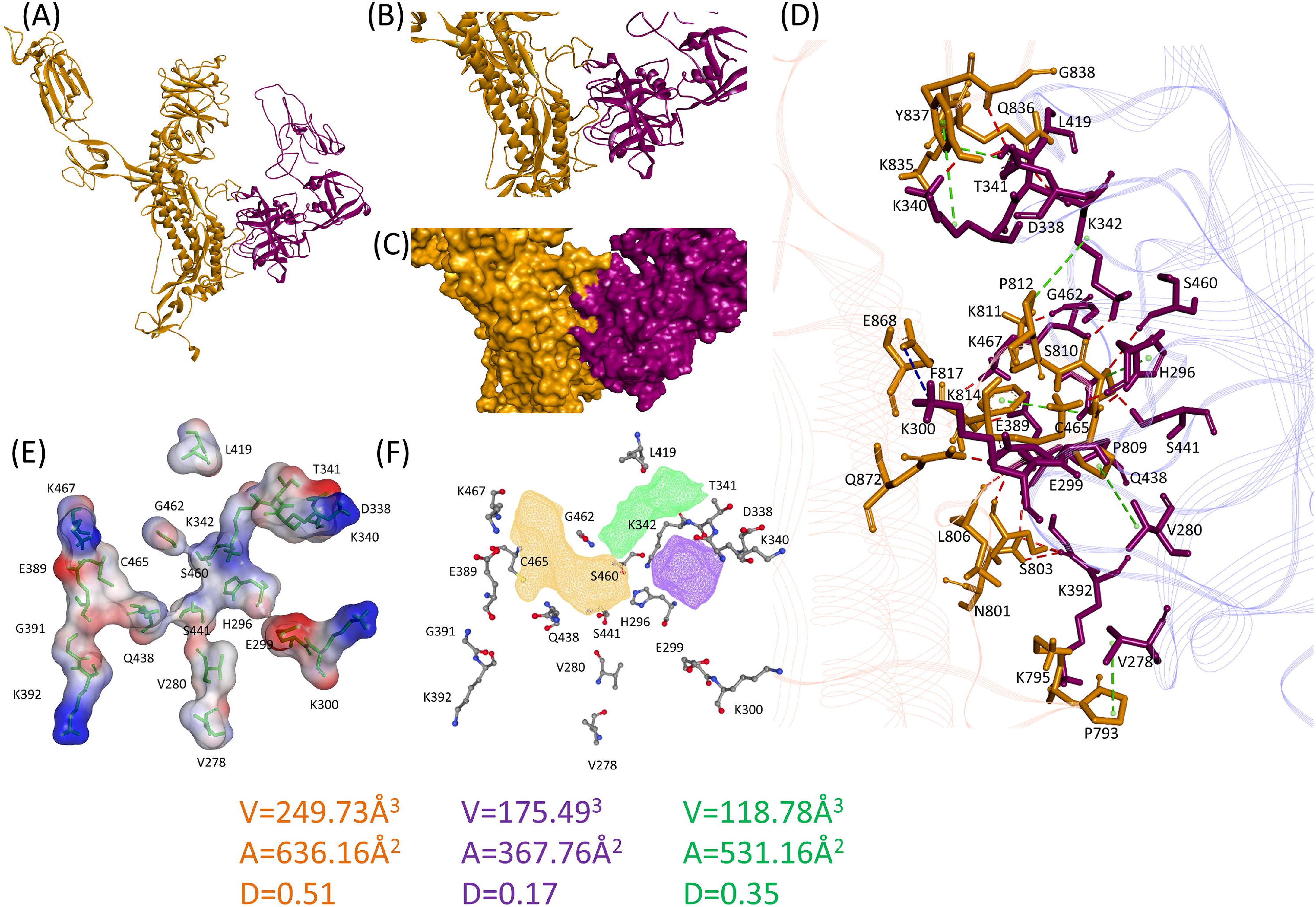

**Table 1:** Comparison of intermolecular interactions between TMPRSS2 and two different cleavage sites of SARS-CoV-2 Spike protein

| Nature and Properties of Interactions | Cleavage Site 1 (Arg685/Ser686) |  | Cleavage Site 2 (Arg815/Ser816) |  |
| --- | --- | --- | --- | --- |
|  | TMPRSS2 | SARS-CoV-2 Spike Protein | TMPRSS2 | SARS-CoV-2 Spike Protein |
| **Intermolecular Hydrogen Bonds** | ARG413 | GLU1072 | HIS296 | PRO809 |
|  | GLU299 | ARG682 | LYS340 | TYR837 |
|  | THR387 | ASP936 | LYS342 | GLN836 |
|  | ARG413 | GLU1072 | LYS342 | SER810 |
|  | TYR414 | GLU1072 | LYS392 | LYS795 |
|  | TYR469 | GLN1071 | GLN438 | SER803 |
|  | HIS296 | ARG682 | GLN438 | LEU806 |
|  | VAL275 | ARG683 | SER441 | SER810 |
|  | GLN276 | ARG683 | LYS467 | GLN935 |
|  | LYS300 | GLN690 | GLY391 | ASN801 |
|  | GLN431 | GLN926 | GLU389 | ASN801 |
|  | ASN433 | SER929 | GLY391 | ASN801 |
|  | ALA466 | LYS933 | GLY391 | SER803 |
|  | GLU388 | SER937 | GLY462 | LYS811 |
|  | LYS390 | THR941 | HIS296 | LYS814 |
|  | GLY391 | SER940 | ASP338 | TYR837 |
|  | ARG413 | GLU1072 | THR341 | GLY838 |
|  | GLU299 | ARG682 | GLU299 | GLN872 |
|  |  |  | GLU389 | GLN935 |
|  |  |  | SER460 | SER810 |
| **Intermolecular Electrostatic Interactions** | Lys300 | GLU654 | LYS300 | GLU868 |
|  | GLU299 | ARG682 |  |  |
|  | GLU299 | ARG682 |  |  |
|  | GLU388 | LYS933 |  |  |
|  | HIS296 | ARG682 |  |  |
|  | ARG413 | GLU1072 |  |  |
| **Intermolecular Hydrophobic Interactions** | VAL278 | ARG683 | HIS296 | SER810 |
|  |  |  | VAL278 | PRO793 |
|  |  |  | VAL280 | PRO809 |
|  |  |  | LYS342 | PRO812 |
|  |  |  | LEU419 | LYS835 |
|  |  |  | CYS465 | PHE817 |
|  |  |  | LYS340 | TYR837 |
| **Charged-Charged** | 9 |  | 11 |  |
| **Charged –Polar** | 21 |  | 32 |  |
| **Charged-Apolar** | 17 |  | 27 |  |
| **Polar-Polar** | 12 |  | 11 |  |
| **Polar-Apolar** | 16 |  | 28 |  |
| **Apolar-Apolar** | 3 |  | 13 |  |
| **Binding Affinity** | -9.8 Kcal/mol |  | -13.8 Kcal/mol |  |
| **Kd at 25°C** | 6.8 X 10<sup>-8</sup> |  | 7.1 X 10<sup>-11</sup> |  |
| **Kd at 37°C** | 1.3 X 10<sup>-7</sup> |  | 1.8 X 10<sup>-10</sup> |  |

## Discussion

Human TMPRSS2 is a 70kDa protein, a member of large superfamily of serine protease, mainly expressed in prostate, colon, stomach, and salivary gland (Vaarala et al., 2001). In prostate gland its expression is regulated by androgens and found overexpressed in prostate carcinoma (Afar et al., 2001). Physiologically, the protein is important in the functioning of epithelial sodium transport (Donaldson et al., 2002) and angiogenesis (Aimes et al., 2003). In addition, TMRPSS2 importance has been demonstrated in relation to the entry of influenza virus (Böttcher et al., 2006), SARS-CoV (Shulla et al., 2011), parainfluenza virus (Abe et al., 2013), MERS-CoV (Shirato et al., 2013) and SARS-CoV-2 (Hoffman et al., 2020). Cleavage sites of SARS-CoV-2 spike protein for TMPRSS2 action have been mapped, but the complex structure of SARS-CoV-2 spike protein and TMPRSS2 has not been resolved. An investigative flank of the unpublished study (DOI: 10.1101/2020.02.08.926006) attempted to address the same issue, however, focusing on the development of peptidyl analogue targeting merely catalytic triad of TMPRSS2 using partial model of the molecule. Additionally, details regarding the interaction between the viral spike protein and TMPRSS2 have not been resolved at the residual level and/or for both cleavage sites.

We have constructed full length model of TMPRSS2 showing distinct localization of all three functional domains. The C-terminal peptidase S1 domain is expectedly involved in the interaction with SARS-CoV-2 spike protein. Both substrate binding and catalytic sites residues of TMPRSS2 interact with the cleavage sites and/or immediate flanking residues of SARS-CoV-2 spike protein. Involvement of the neighbouring amino acids to the active site of TMPRSS2 and cleavage sites of SARS-CoV-2 provides important clues for the design of targeted inhibitors and/or peptidyl disruptors. Several protease inhibitors have been proposed by means of virtual screening (Shah et al., 2020) and have shown efficacy against SARS-CoV-2 infection (Hoffman et al., 2020). The findings of the present study in relation to the diversity of the nature of intermolecular interactions and biophysciochemical properties of entailing amino acids may in turn facilitate structure based drug designing for the more efficient peptidyl antagonists against COVID-19. Peptidyl inhibitors have shown to efficiently inhibit EBNA1 dimerization (Hussain, 2013), protein-protein interactions of coiled-coiled transcription factors like Bcl-2 proteins, MDM2/MDMX, HIVgp41 (Aragi and Keating, 2016) and human thymidylate synthase (Cardinale et al., 2011). In addition, the present study further points to the key residues for the subsequent investigations like site directed mutagenesis and peptide array studies to discern importance of potentially other residues in the priming of the viral spike protein. Recently, we have reported that allelic variants in the human ACE2 receptor flanking the key interacting amino acids may hamper its attachment with SARS-CoV-2 spike protein (Hussain et al., 2020). Therefore, the present study could also be advanced in relation to explore the effect of natural polymorphism found in human TMPRSS2 on priming of SARS-CoV-2 spike protein.

The authors declare that they have no conflict of interest

